# Long-read genome sequence and assembly of *Leptopilina boulardi*: a specialist *Drosophila* parasitoid

**DOI:** 10.1101/284679

**Authors:** Shagufta Khan, Divya Tej Sowpati, Rakesh K Mishra

## Abstract

**Background:** *Leptopilina boulardi* is a specialist parasitoid belonging to the order Hymenoptera, which attacks the larval stages of *Drosophila*. The *Leptopilina* genus has enormous value in the biological control of pests as well as in understanding several aspects of host-parasitoid biology. However, none of the members of Figitidae family has their genomes sequenced. In order to improve the understanding of the parasitoid wasps by generating genomic resources, we sequenced the whole genome of *L. boulardi*.

**Findings:** Here, we report a high-quality genome of *L. boulardi*, assembled from 70Gb of Illumina reads and 10.5Gb of PacBio reads, forming a total coverage of 230X. The 375Mb draft genome has an N50 of 275Kb with 6315 scaffolds >500bp, and encompasses >95% complete BUSCOs. The GC% of the genome is 28.26%, and RepeatMasker identified 868105 repeat elements covering 43.9% of the assembly. A total of 25259 protein-coding genes were predicted using a combination of *ab-initio* and RNA-Seq based methods, with an average gene size of 3.9Kb. 78.11% of the predicted genes could be annotated with at least one function.

**Conclusion:** Our study provides a highly reliable assembly of this parasitoid wasp, which will be a valuable resource to researchers studying parasitoids. In particular, it can help delineate the host-parasitoid mechanisms that are part of the *Drosophila* – *Leptopilina* model system.

## Data Description

Parasitoids are organisms that have a non-mutualistic association with their hosts. Nearly 20% of the identified insects are known to be parasitoids, the vast majority of which are parasitoid wasps belonging to the order Hymenoptera [1]. Parasitoid wasps are classified into two categories based on their host preference – generalists and specialists. Generalists can infect a wide range of species whereas specialists parasitize one or two host species. *Leptopilina boulardi* (NCBI taxonomy ID: 63433) is a solitary endoparasitoid wasp from the Figitidae family in the Hymenoptera order (Fig 1). It is a cosmopolitan species, ubiquitously found in the Mediterranean and intertropical environments, having its origin from Africa [2]. *L. boulardi* succeeds in parasitizing *D. melanogaster* and *D. simulans* at second-to early third-instar larval stages and hence, is referred to as a specialist [3].

**Figure 1:**
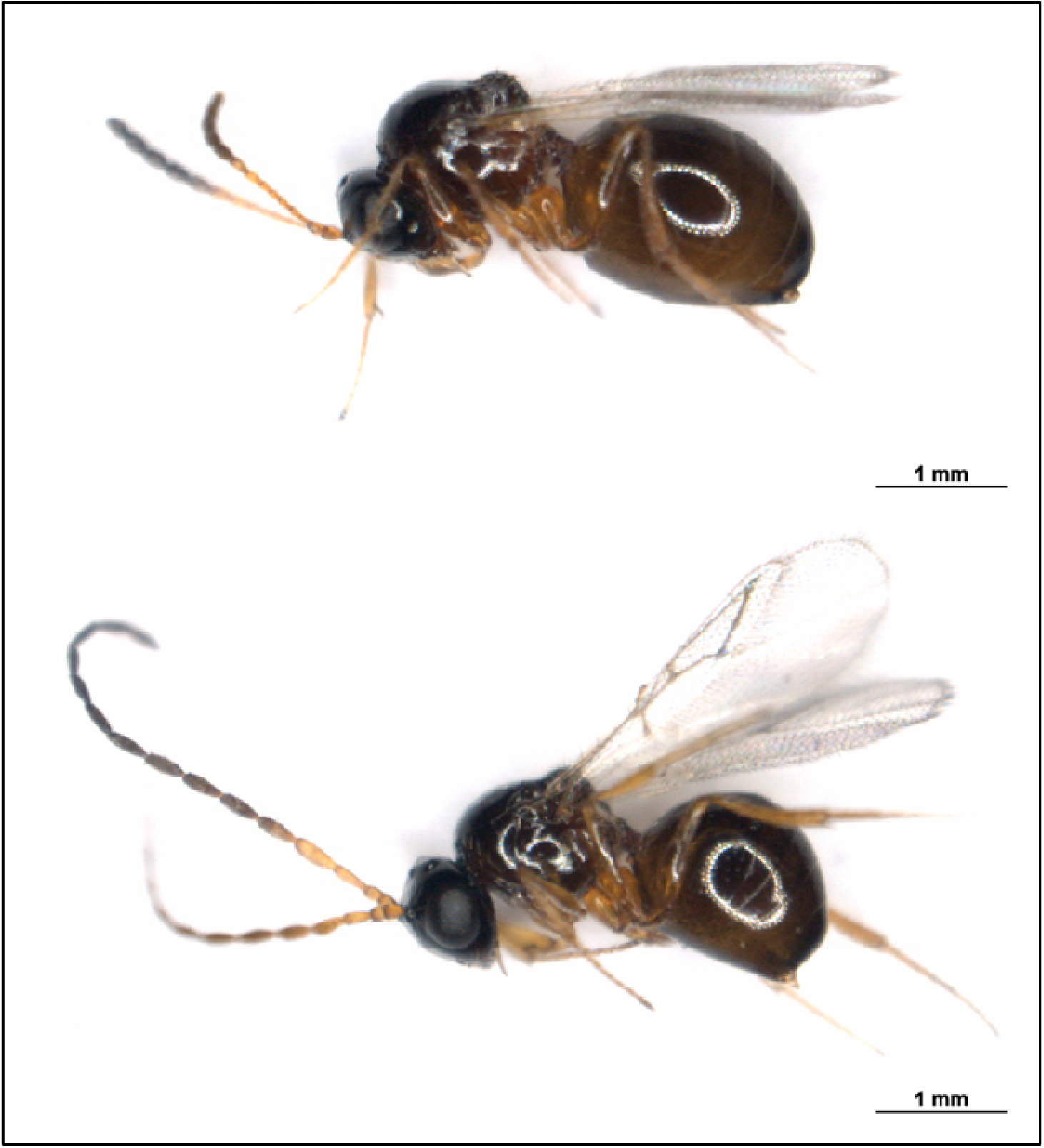
*Leptopilina boulardi* (Lb17 strain) – adult female (top) and male (bottom)

Similar to other Drosophilid parasitoids, *L. boulardi* has a haplodiploid sex-determination system; the unfertilized eggs and fertilized eggs develop into haploid males and diploid females respectively [4]. The females of this figitid species are endoparasitic koinobionts, i.e., they lay eggs inside the host’s larva, allowing the host to grow and feed without rapidly killing it [1]. During oviposition, the wasps co-inject virulence factors like venom proteins, Virus-like Particles (VLPs) into the larval hemolymph that help in evading the host’s immune responses [5-7]. After hatching inside the host hemocoel, the wasp larva histolyzes the host tissues gradually. Subsequently, the endoparasitoid transitions into an ectoparasitoid and consumes the host entirely while residing inside the host puparium until emergence. The entire life cycle takes 21-22 days at 25°C [2, 8]. Alternatively, the host elicits an immune response leading to the encapsulation followed by the death of the parasitoid and emergence of the host [9, 10]. The virulence of the parasitoid wasps varies with the strain and species, the genetic basis of which remains unclear.

Apart from the potential use of Figitidae parasitoids in biological control of pests, *Drosophila* – *Leptopilina* system has been intensively studied to understand various aspects of the host-parasitoid biology like coevolutionary dynamics, behavioral ecology, innate-immune responses, and superparasitism [3, 11-13]. The cytogenetic and karyotypic analysis has revealed interesting features about the genome size and chromosome number of numerous parasitoid species [14]. However, except for the mitochondrial genome of *L. boulardi* [15], none of the genomes of the members of the estimated 24,000 species [16] in the Figitidae family has been sequenced, greatly limiting the scope of the field. Here, we provide the first complete reference genome of *L. boulardi*, a Figitid parasitoid, for a better understanding of this emerging model system.

### Sample Collection

*L. boulardi* (Lb17 strain), kindly provided by S. Govind (Biology Department, The City College of the City University of New York), was reared on *D. melanogaster* (Canton-S strain) as described earlier [10]. Briefly, 50-60 young flies were allowed to lay eggs for 24 hours at 25°C in vials containing standard yeast/corn-flour/sugar/agar medium. Subsequently, the host larvae were exposed to 6-8 male and female wasps, respectively, 48 hours after the initiation of egg lay. The culture conditions were maintained at 25°C and LD 12:12. The wasps (2 days old) were collected, flash-frozen in liquid nitrogen, and stored at -80°C until further use.

### Genomic DNA preparation

For whole genome sequencing on Illumina HiSeq 2500 platform (Table 1), the genomic DNA was extracted as follows: 100 mg of wasps were ground into a fine powder in liquid nitrogen and kept for lysis at 55°C in SNET buffer (400 mM NaCl, 1% SDS, 20 mM Tris-HCl pH8.0, 5 mM EDTA pH 8.0 and 2 mg/ml Proteinase K) with gentle rotation at 10 rpm overnight. Next day, after RNase A (100 μg/ml) digestion, Phenol:Chloroform:Isoamyl Alcohol extraction was performed followed by Ethanol precipitation. The pellet was resuspended in 1X Tris-EDTA buffer (pH 8.0).

**Table 1:**
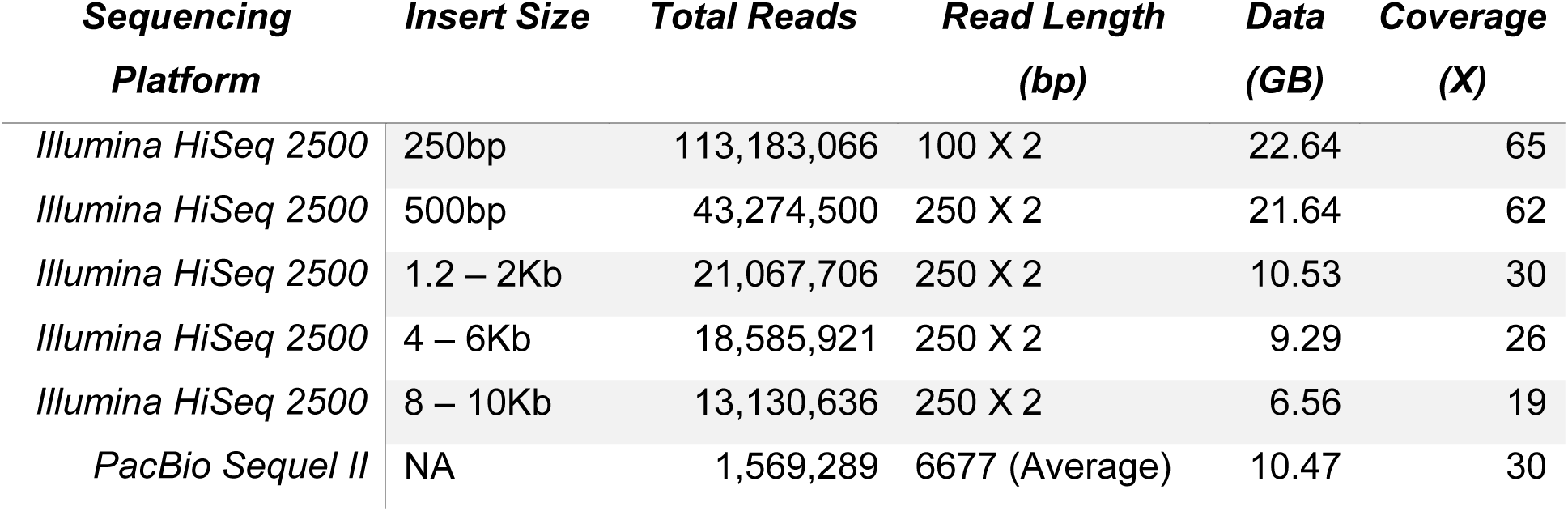
Details of the sequencing data generated for the genome assembly of *L. boulardi*.

For long-read sequencing on PacBio Sequel II platform, the genomic DNA preparation was done from 200 mg wasps using the protocol described earlier [17] with the following additional steps: Proteinase K digestion for 30 minutes at 50°C after lysis, RNase A digestion for 10-15 minutes at RT (1 μl per 100 μl of 100 mg/ml) after the centrifugation step of contaminant precipitation with potassium acetate and a single round of Phenol:Chloroform:Isoamyl Alcohol (25:24:1, v/v) (Cat. No. 15593031) phase separation before genomic DNA purification using Agencourts AMPure XP beads (Item No. A63880).

### Hybrid assembly of short and long reads

Cytogenetic analysis has estimated the genome size of *L. boulardi* to be around 360Mb [14]. We used JellyFish [18] to determine the genome size of *L. boulardi* to be 398Mb. Assembly of the reads was done using a hybrid assembler, MaSuRCA [19]. MaSuRCA uses both short Illumina reads and long PacBio reads to generate error-corrected super reads, which are further assembled into contigs. It then uses mate-pair information from short read libraries to scaffold the contigs. Using the 5 short read libraries of ∼200X coverage (70.66GB data) and PacBio reads of ∼30X coverage (10.5GB data), MaSuRCA produced an assembly of 375Mb, made of 6341 scaffolds with an N50 of 275Kb (Table 2). The largest scaffold was 2.4Mb long, and 50% of the assembly was covered by 380 largest scaffolds (L50). GapFiller [20] was used to fill N’s in the assembly. After 10 iterations, 206Kb out of 1.4Mb of N’s could be filled using GapFiller. From this assembly, all scaffolds shorter than 500bp were removed, leaving a total of 6315 scaffolds. This version was used for all further analyses.

**Table 2:**
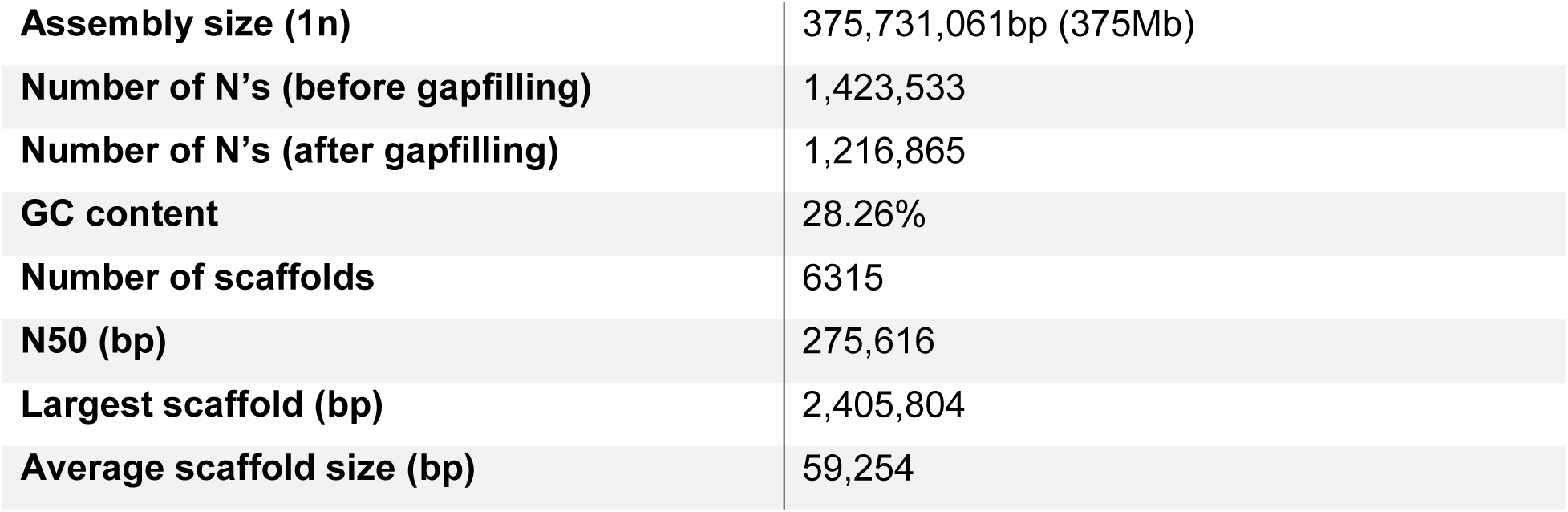
Summary statistics of the assembled *L. boulardi* genome.

### Assessment of genome completeness

The quality of the genome assembly was measured using two approaches. First, we aligned the paired end reads of the 250bp library to the assembly using bowtie2 [21]. 94.64% of the reads could be mapped back, with 92.32% reads mapped in proper pairs. Next, we used BUSCO v3 [22] to look for the number of single-copy orthologs in the assembly. Out of the 978 BUSCOs in the metazoan dataset, 943 (96.5%) complete BUSCOs were detected in the assembly (Table 3). We also performed BUSCO analysis with the Arthropoda (1066 BUSCOs) and Insecta (1658 BUSCOs) datasets, and could identify 97% and 95.7% complete BUSCOs in our assembly respectively (Table 3). Both the results indicated that the generated assembly was nearly complete, with a good representation of the gene repertoire.

**Table 3:**
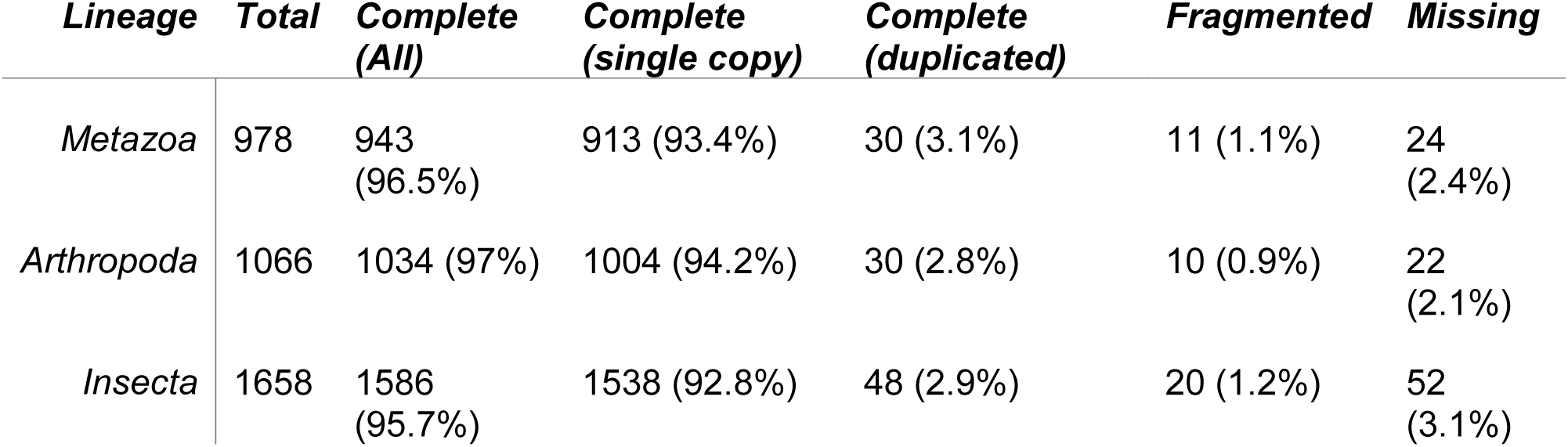
BUSCO analysis of the *L. boulardi* genome.

| **Lineage** | **Total** | **Complete**<br>**(All)** | **Complete**<br>**(single copy)** | **Complete**<br>**(duplicated)** | **Fragmented** | **Missing** |
| --- | --- | --- | --- | --- | --- | --- |
| *Metazoa* | 978 | 943<br>(96.5%) | 913 (93.4%) | 30 (3.1%) | 11 (1.1%) | 24<br>(2.4%) |
| *Arthropoda* | 1066 | 1034 (97%) | 1004 (94.2%) | 30 (2.8%) | 10 (0.9%) | 22<br>(2.1%) |
| *Insecta* | 1658 | 1586<br>(95.7%) | 1538 (92.8%) | 48 (2.9%) | 20 (1.2%) | 52<br>(3.1%) |

### Identification of repeat elements

To identify repeat elements in the *L. boulardi* assembly, we first used RepeatModeler with RepeatScout [23] and TRF [24] to generate a custom repeat library. The output of RepeatModeler was provided to RepeatMasker [25], along with the RepBase library [26], to search for various repeat elements in the assembly. Table 4 summarizes the number of repeat elements identified as well as their respective types. A total of 868105 repeat elements could be identified using RepeatMasker, covering almost 165Mb (43.88%) of the genome. We further used PERF [27] to identify simple sequence repeats of >=12bp length. PERF reported a total of 853,624 SSRs covering 12.24Mb (3.26%) of the genome (Table 5). Hexamers were the most abundant SSRs (40.1%) in *L. boulardi*, followed by pentamers (15.8%) and monomers (14.3%).

**Table 4:** Summary of repeat elements identified by RepeatMasker in the *L. boulardi* genome.

| *Repeat Type* | *Number of Elements* | *Length in bp* | *% Genome Covered* |
| --- | --- | --- | --- |
| *SINEs* | 3721 | 1,651,220 | 0.44 |
| *LINEs* | 10573 | 5,613,129 | 1.49 |
| *LTR elements* | 12312 | 9,512,954 | 2.53 |
| *DNA elements* | 105817 | 31,232,845 | 8.31 |
| *Unclassified interspersed elements* | 382214 | 102,924,940 | 27.39 |
| *Small RNA* | 186 | 137,204 | 0.04 |
| *Satellites* | 2442 | 1,028,732 | 0.27 |
| *Simple repeats* | 251669 | 11,461,332 | 3.05 |
| *Low complexity* | 46977 | 2,473,942 | 0.66 |

**Table 5:**
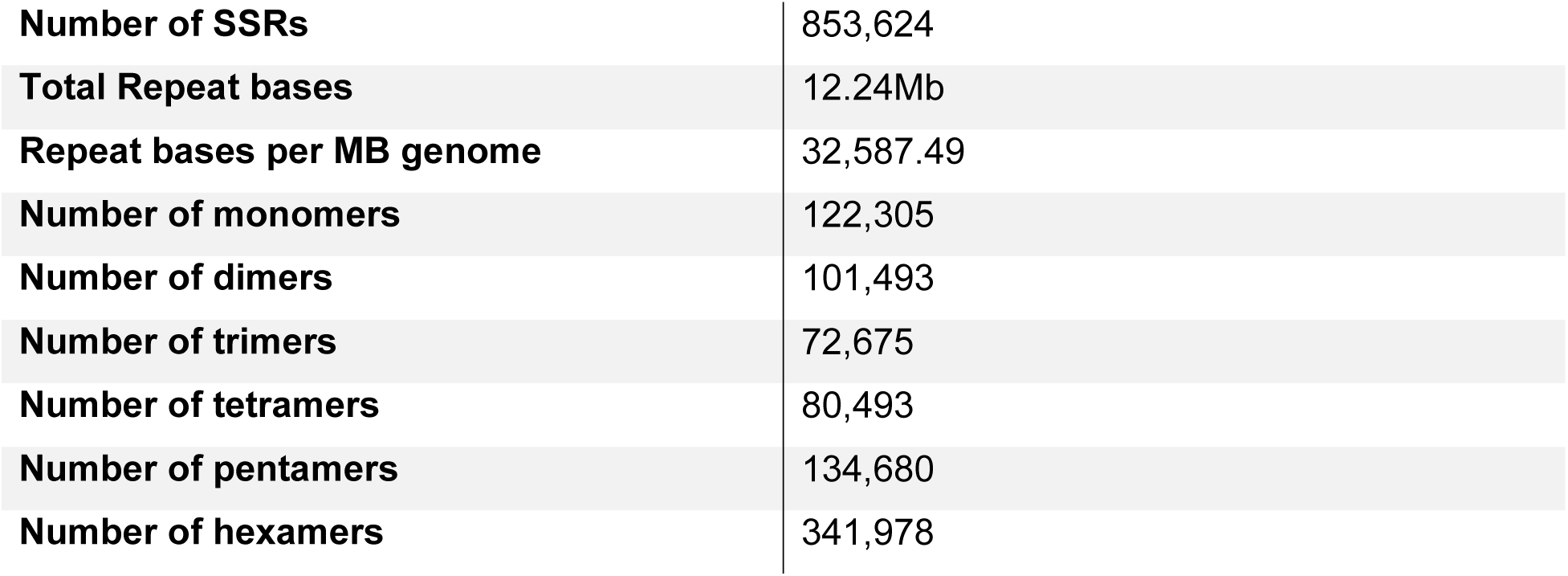
Details of Simple Sequence Repeats identified by PERF in the *L. boulardi* genome.

### Gene prediction

Coding regions in the assembled genome of *L. boulardi* were predicted using two different approaches: RNA-seq based prediction and *ab initio* prediction. For RNA-seq based approach, available paired-end data generated from the transcriptome of *L. boulardi* (SRR559222) was mapped to the assembly using STAR [28]. The BAM file containing uniquely-mapped read pairs (72% of total reads) was used to construct high quality transcripts using Cufflinks [29]. The same BAM file was submitted for RNA-seq based *ab initio* prediction using BRAKER [30]. BRAKER uses the RNA-seq data to generate initial gene structures using GeneMark-ET [31], and further uses AUGUSTUS [32] to predict genes based on the generated gene structures. In addition to BRAKER, two other *ab initio* prediction tools were used: GlimmerHMM [33] and SNAP. The number of predicted genes using each method is outlined in Table 6. Using the gene sets generated from various methods, a final non-redundant set of 25259 genes was derived using Evidence Modeler [34] (Table 6). The average gene size in the final gene set is ∼3.9Kb. A protein FASTA file was derived using this gene set, which was used for functional annotation.

**Table 6:**
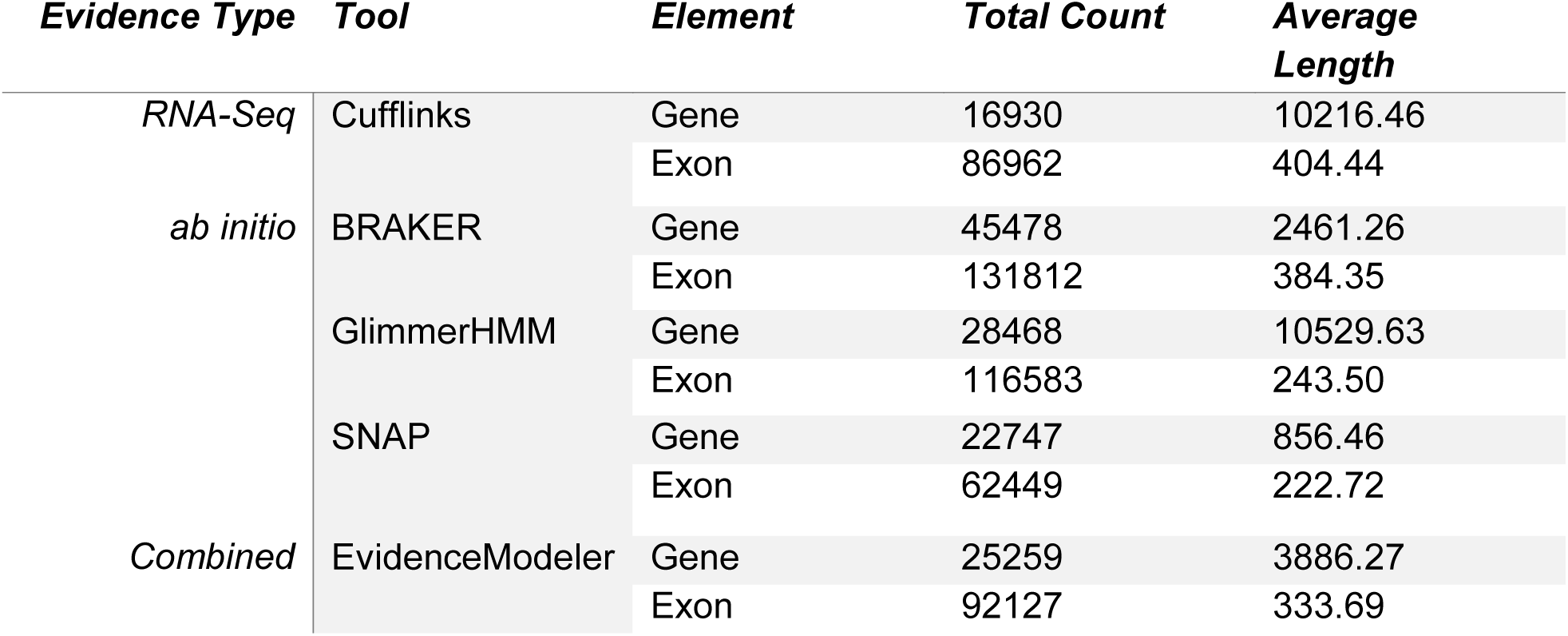
Prediction of genes in *L. boulardi*: summary of various methods.

| *Evidence Type* | *Tool* | *Element* | *Total Count* | *Average Length* |
| --- | --- | --- | --- | --- |
| RNA-Seq | Cufflinks | Gene | 16930 | 10216.46 |
|  |  | Exon | 86962 | 404.44 |
| *ab initio* | BRAKER | Gene | 45478 | 2461.26 |
|  |  | Exon | 131812 | 384.35 |
|  | GlimmerHMM | Gene | 28468 | 10529.63 |
|  |  | Exon | 116583 | 243.50 |
|  | SNAP | Gene | 22747 | 856.46 |
|  |  | Exon | 62449 | 222.72 |
| Combined | EvidenceModeler | Gene | 25259 | 3886.27 |
|  |  | Exon | 92127 | 333.69 |

### Gene annotation

The functional annotation of predicted proteins was done using homology-based approach. InterProScan v5 [35] was used to search for homology of protein sequences against various databases such as Pfam, PROSITE, and Gene3D. 12,449 out of 25,259 (49.2%) proteins could be annotated using Pfam, while 9346 and 10952 proteins showed a match in PROSITE and Gene3D databases respectively (Table 7). The gene ontology terms associated with the proteins were retrieved using the InterPro ID assigned to various proteins. A total of 19731 proteins (78.11%) could be annotated using at least one database.

**Table 7:**
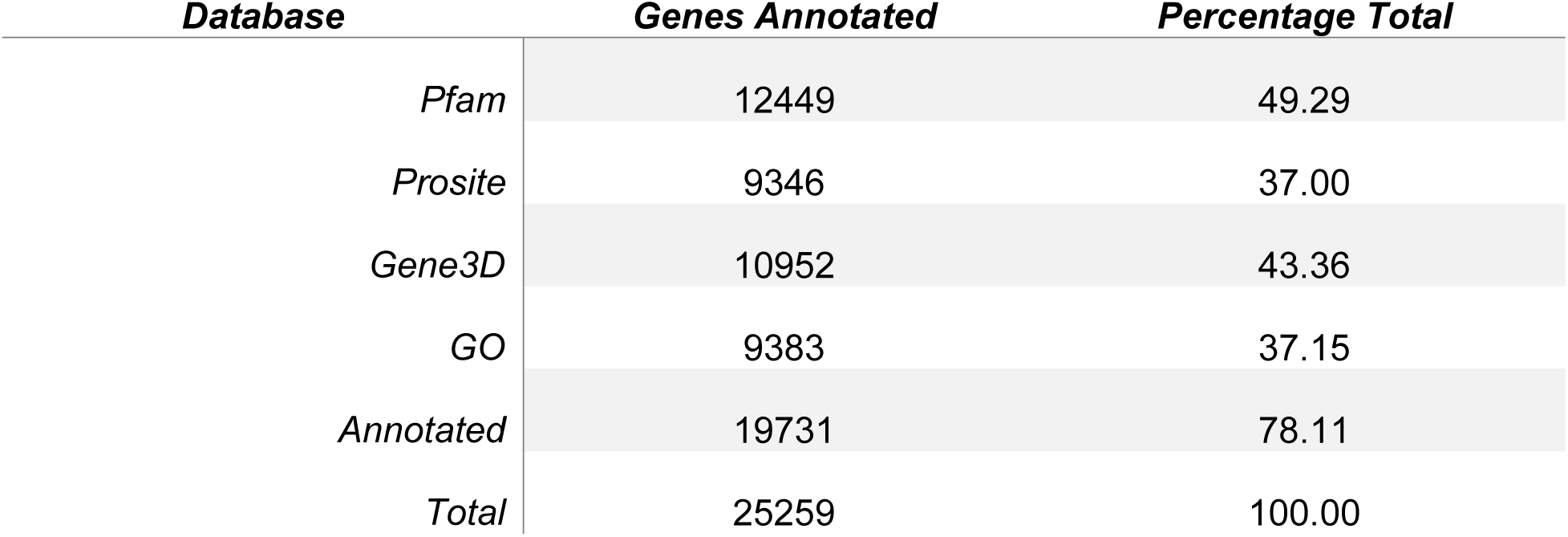
Gene Annotation of the predicted genes in *L. boulardi*.

## Conclusions

Our study reports a high-quality genome of the specialist parasitoid wasp *Leptopilina boulardi.* BUSCO analysis showed almost a complete coverage of the core gene repertoire. A total of 25,259 protein-coding genes were predicted, out of which 19731 could be annotated using known protein signatures. This genome thus provides a valuable resource to researchers studying parasitoids, and can help shed some light on the mechanisms of host-parasitoid interactions, and understanding the immune response mechanisms in insects. Being the first complete genome from the Figitidae family, the genome sequence of *L. boulardi* will also be a key element in understanding the evolution of parasitism in Figitids.

## Availability of Data

The raw reads generated on the Illumina and PacBio platforms will be available on the Sequence Read Archive (SRA) of NCBI. The assembled scaffolds, predicted gene and protein sequences will be available from the genome repository of NCBI.

## Acknowledgements

We thank Shubha Govind, The City College of the City University of New York, for providing us the Lb17 strain of *L. boulardi*. We acknowledge Indira Paddibhatla for introducing the *Drosophila-Leptopilina* system to our lab. Athira Rajeev is acknowledged for the initial quality control and processing of Illumina data.

This work is supported by the Genesis project of the Council of Scientific and Industrial Research, India (BSC0121). SK is a recipient of the DST-INSPIRE Research Fellowship.

